# Decoupling the variances of heterosis and inbreeding effects is evidenced in yeast’s life-history and proteomic traits

**DOI:** 10.1101/362418

**Authors:** M. Petrizzelli, D. de Vienne, C. Dillmann

**Author notes:** To whom correspondence should be addressed: UMR Génétique Quantitative et Évolution - Le Moulon, Ferme du Moulon, 91190 Gif-sur-Yvette, France.

## Abstract

Heterosis (hybrid vigor) and inbreeding depression, commonly considered as corollary phenomena, could nevertheless be decoupled under certain assumptions according to theoretical population genetics works. In order to explore this issue on real data, we analyzed the components of genetic variation in a population derived from a half-diallel cross between strains from *Saccharomyces cerevisiae* and *S. uvarum*, two related yeast species involved in alcoholic fermentation. A large number of phenotypic traits, either molecular (coming from quantitative proteomics) or related to fermentation and life-history, were measured during alcoholic fermentation. Because the parental strains were included in the design, we were able to distinguish between inbreeding effects, which measures phenotypic differences between inbred and hybrids, and heterosis, which measures phenotypic differences between a specific hybrid and the other hybrids sharing a common parent. The sources of phenotypic variation differed depending on the temperature, indicating the predominance of genotype by environment interactions. Decomposing the total genetic variance into variances of additive (intra- and inter-specific) effects, of inbreeding effects and of heterosis (intra- and inter-specific) effects, we showed that the distribution of variance components defined clear-cut groups of proteins and traits. Moreover, it was possible to cluster fermentation and life-history traits into most proteomic groups. Within groups, we observed positive, negative or null correlations between the variances of heterosis and inbreeding effects. To our knowledge, such a decoupling had never been experimentally demonstrated. This result suggests that, despite a common evolutionary history of individuals within a species, the different types of traits have been subject to different selective pressures.

---

Heterosis, or hybrid vigor, refers to the common superiority of hybrids over their parents for quantitative traits. This phenomenon has been observed for virtually any quantitative trait, from mRNA abundances to fitness, and in a large diversity of species, including microorganisms. For decades it has been extensively studied and exploited for plant and animal breeding, since it affects traits of high economical interest such as biomass, fertility, growth rate, disease resistance etc. (Gowen 1952; Schnable and Springer 2013).

There are three classical, non exclusive genetic models to account for hybrid vigor: dominance, overdominance and epistasis. In the dominance model, the hybrid superiority results from the masking of the deleterious alleles of one parent by the non deleterious ones of the other parent (Davenport 1908). In the overdominance model, the hybrid superiority is due to the advantage *per se* of the heterozygous state at a given locus (Hull 1946). Actually, more common is pseudo-overdominance, which is due to dominance at two loci linked in repulsion, *e*.*g*. in maize (Graham *et al.* 1997; Lariepe *et al.* 2012) or yeast (Martì-Raga *et al.* 2017). Lastly, the epistasis model postulates favorable intergenic interactions created in the hybrids (Powers 1944). In particular, “less-than-additive” (antagonistic) epistasis, which is quite common in plant and animal species (Redden 1991; Shao *et al.* 2008) can account for best-parent heterosis (Fievet *et al.* 2010). In this last paper, it is theoretically shown that epistasis can result in best-parent heterosis even if there is no dominance at any locus. The respective parts of the various genetics effects in heterosis depends on the trait, the species and the genetic material (Xiao *et al.* 1995; Huang *et al.* 2016; Seymour *et al.* 2016). Altogether, heterosis appears to be a pervasive phenomenon, accounted for by the common non-linearity of the genotype-phenotype map (Wright 1934; Omholt *et al.* 2000; Fiévet *et al.* 2018).

Because heterosis is associated with heterozygosity, heterosis for life-history traits is associated with genetic load: the average population fitness can never exceed the maximum fitness. Genetic load drives the evolution of sexual reproduction, of mating systems as well as the fate of small populations. Indeed, high levels of homozygosity in outcrossing species is generally associated with decreased growth rate, survival or fertility (discussed in Charlesworth and Willis (2009)). In population genetics, inbreeding depression is defined as the fitness of self-fertilized progenies as compared with fitness of outcrossing progenies. In sexual species, the balance between selfing and outcrossing is driven by the genetic load due to inbreeding depression relative to the cost of sexual reproduction (twice as expensive as clonal reproduction): selfing can evolve whenever inbreeding depression is less costly than the sexual reproduction, or after purging deleterious mutations as can arise in small populations (Lande and Schemske 1985). However, heterosis due to less-than-additive epistasis could explain the large number of predominantly (but not fully) selfing species exhibiting a persistent amount of inbreeding depression and heterosis (Charlesworth *et al.* 1991). Considering a metapopulation, Roze and Rousset (2004) defined inbreeding depression as the fitness reduction of selfed progeny relative to outcrossed progeny within populations, and heterosis as the difference between the fitness of the outcrossed progeny within population and the outcrossed progeny over the whole metapopulation. They showed that while selfing reduced both inbreeding depression and heterosis, inbreeding depression decreased and heterosis increased with the degree of subdivision of the metapopulation. Hence, from a population genetics point of view, heterosis is expected even in predominantly selfing species.

In a breeding perspective, the pioneer work of Shull (1908) in maize predicted that given the large amounts of heterosis within the species, the best way to maximize yield was to create inbreds from existing population varieties in order to seek for the best hybrid combinations. Diallel designs were popularized as the most comprehensive designs for estimating genetic effects, predicting hybrid values and generating breeding populations to be used as basis for selection and development of elite varieties (i.e. Hallauer and Filho (1988)). The simplest and most popular analytic decomposition of genetic effects in diallel designs is that of Griffing (1956), in which the mean phenotypic value, *y_ij_*, of the cross between lines *i* and *j* is modeled as:

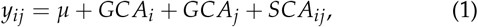

where *μ* is the mean phenotypic value of the population, *GCAi* (resp. *GCAj*) is the *General Combining Ability* of line *i* (resp. *j*), *i*.*e*. the average performance of line *i* (resp. *j*) in hybrid combinations expressed as a deviation from the mean value of all crosses, and *SCA_ij_* is the *Specific Combining Ability* of hybrid *i ×j*. It is defined as the difference between the mean phenotypic value of the progeny and the sum of the combining abilities of the parental lines (Sprague and Tatum 1942). Therefore, superior individuals can be selected from their GCA and/or SCA. Numerous extensions of the Griffing’s model have been proposed to extract other effects, such as maternal and paternal effects or sex-linked variations (Cockerham and Weir 1977; Bulmer 1980; Zhu and Weir 1996; Greenberg *et al.* 2010). In many crop species, combining ability groups have been identified, with lines from the same group characterized by high specific combining ability with other groups (Hallauer *et al.* 1988). Generally, combining ability groups are redundant with population structure within a species (Melchinger and Gumber 1998; Ramya *et al.* 2018), which is consistent with the population genetics predictions of Roze and Rousset (2004).

When parental lines are included in the analysis, *GCA* and *SCA* effects can be decomposed in more suitable genetic effects. Indeed, the value of a particular hybrid can be compared either to the average value of its inbred parents, or to the average value of the other hybrids sharing either parent. Heterosis can be split into *average heterosis* (average difference between inbreds and outbreds), *variety heterosis* (average difference between one inbred parent and all crosses sharing the same parents), and *specific heterosis* (difference between the hybrid and all hybrids sharing at least one parent) (Eberhart and Gardner 1966). A modern version of this model have been proposed by Lenarcic *et al.* (2012) along with a Bayesian framework to estimate the genetic effects.

In this work, we study a half-diallel design with diagonal constructed from the crosses between 11 yeast strains belonging to two close species, *Saccharomyces cerevisiae* and *S. uvarum*. The design included both intra- and inter-specific crosses. Two categories of phenotypic traits were considered: (i) protein abundances measured at one time point of alcoholic fermentation (Blein-Nicolas *et al.* 2013, 2015); (ii) a set of fermentation traits measured during and/or at the end of fermentation, which were divided into kinetic parameters, basic enological parameters, aromas and life-history traits (da Silva *et al.* 2015). All traits were independently measured at two temperatures.

We propose a decomposition of the genetic effects based on Lenarcic *et al.* (2012) that takes into account the presence of two species in the diallel design and that distinguishes heterosis and inbreeding effects. We could characterize every trait by the set of its variance components and we could clearly cluster the traits from this criterion, which suggests that traits sharing a similar pattern of variance components could share common life-history. We were able to assign each fermentation trait to one group of protein traits, which shows that integrated phenotypes and proteins can share similar life-history. Finally, our results show a poor correlation between the variances of heterosis and inbreeding effects within groups. This confirms the importance of epistatic interactions in determining the components of phenotypic variation both within and between close species. Altogether, our results suggest that despite a common demographic history of individuals within a species, the genetic variance components of the traits can be used to trace back other trait-specific evolutionary pressures, like selection.

## Materials and Methods

### Materials

The genetic material of the experimental design consisted in 7 strains of *S. cerevisiae* and 4 strains of *S. uvarum* associated to various food-processes (enology, brewery, cider fermentation and distillery) or isolated from natural environment (oak exudates). These strains, called W1, D1, D2, E2, E3, E4, E5 for *S. cerevisiae* and U1, U2, U3, U4 for *S. uvarum* could not be used as such as parents of a diallel design because they were suspected to be heterozygous at many loci. Monosporic clones were isolated from each of these strains using a micromanipulator (Singer MSM Manual; Singer Instrument, Somerset, United Kingdom), as indicated in da Silva *et al.* (2015). All strains but D2 were homothallic (HO/HO), therefore fully homozygous diploid strains were spontaneously obtained by fusion of opposite mating type cells. For D2 (ho/ho), the isolated haploid meiospore were diploidized via transient expression of the HO endonuclease (Albertin *et al.* 2009). The derived fully homozygous and diploid strains were used as the parental strains of a half-diallel design with diagonal, *i.e.* including the inbred lines. The parental lines were selfed and pairwise crossed, which resulted in a total of 66 strains: 11 inbred lines, 27 intra-specific hybrids (21 for *S. cerevisiae*, noted *S. c.*, and 6 for *S. uvarum*, noted *S. u.*) and 28 inter-specific (noted *S. u.× S. c*). For each hybrid construction, parental strains of opposite mating type were put in contact for 2 to 6 hours in YPD medium at room temperature, and then plated on YPD-agar containing the appropriate antibiotics. The nuclear and mitochondrial stability of the hybrids was checked after recurrent cultures on YPD-agar corresponding to ≈ 80 generations (see details in Albertin *et al.* (2013a)). In addition, for each of the 28 interspecific hybrids, both parental sets of more than 600 proteins were detected in a proteomic approach Blein-Nicolas *et al.* (2015), with no evidence of hybrid instability.

The 66 strains were grown in triplicate in fermentors at two temperatures, 26° and 18°, in a medium close to enological conditions (Sauvignon blanc grape juice) (da Silva *et al.* 2015). From a total of 396 alcoholic fermentations (66 strains × 2 temperatures × 3 replicas), 31 failed due to poor fermenting abilities of some strains. The design was implemented considering a block as two sets of 27 fermentations (26 plus a control without yeast to check for contamination), one carried out at 26° and the other at 18°. The distribution of the strains in the block design was randomized to minimize the residual variance of the estimators of the strain and temperature effects, as described in Albertin *et al.* (2013b).

For each alcoholic fermentation, two types of phenotypic traits were measured or estimated from sophisticated data adjustment models: 35 fermentation traits and 615 protein abundances.

The fermentation traits were classified into four categories (da Silva *et al.* 2015):

- *Kinetics parameters*, computed from the CO_2_ release curve modeled as a Weibull function fitted on CO_2_ release quantification monitored by weight loss of bioreactors: the fermentation lag-phase, *t-lag* (h); the time to reach the inflection point out of the fermentation lag-phase, *t-Vmax* (h); the fermentation time at which 45 gL^−1^ and 75 gL^−1^ of CO_2_ was released, out of the fermentation lag-phase, *t-45* (h) and *t-75* (h) respectively; the time between *t-lag* and the time at which the CO_2_ emission rate became less than, or equal to, 0.05gL^−1^h^−1^, *AFtime* (h); the maximum CO_2_ release rate, *V*_max_ (gL^−1^ *h^−^*^1^); and the total amount of CO_2_ released at the end of the fermentation, CO_2_max (gL^−1^).
- *Life history traits*, estimated and computed from the cell concentration curves over time, modeled from population growth, cell size and viability quantified by flow cytometry analysis: the growth lag-phase, *t-N*_0_(*h*); the carrying capacity, *K* (log[cells/mL]); the time at which the carrying capacity was reached, *t-N_max_* (h); the intrinsic growth rate, *r* (log[cell division/mL/h]); the maximum value of the estimated CO_2_ production rate divided by the estimated cell concentration, *J_max_* (gh^−1^10^−8^cell^−1^); the average cell size at *t-N_max_*, *Size-t-N_max_* (*μ*m); the percentage of living cells at *t-N_max_*, *Viability-t-N_max_* (%); and the percentage of living cells at *t-75*, *Viability-t-75* (%).
- *Basic enological parameters*, quantified at the end of fermentation: *Residual Sugar* (gL^−1^); *Ethanol* (%vol); the ratio between the amount of metabolized sugar and the amount of released ethanol, *Sugar.Ethanol.Yield* (gL^−1^%vol^−1^); *Acetic acid* (gL^−1^ of H_2_SO_4_); *Total SO*_2_ (mgL^−1^) and *Free SO*_2_ (mgL^−1^).
- *Aromatic traits*, mainly volatile compounds measured at the end of alcoholic fermentation by GC-MS: two higher alcohols (*Phenyl-2-ethanol* and *Hexanol*, mgL^−1^); seven esters (*Phenyl-2-ethanol acetate*, *Isoamyl acetate*, *Ethyl-propanoate*, *Ethyl-butanoate*, *Ethyl-hexanoate*, *Ethyl-octanoate* and *Ethyl-decanoate*, mgL^−1^); three medium chain fatty acids (*Hexanoic acid*, *Octanoic acid* and *Decanoic acid*, mgL^−1^); one thiol *4-methyl-4-mercaptopentan-2-one*, *X4MMP*(mgL^−1^) and the acetylation rate of higher alcohols, *Acetate ratio*.

For proteomic analyses the samples were harvested at 40 % of CO_2_ release, corresponding to the maximum rate of CO_2_ release. Protein abundances were measured by LC-MS/MS techniques from both shared and proteotypic peptides relying on original Bayesian developments (Blein-Nicolas *et al.* 2012). A total of 615 proteins were quantified in more than 122 strains × temperature combinations as explained in details in Blein-Nicolas *et al.* (2015).

Cross-referencing MIPS micro-organism protein classification (Ruepp *et al.* 2004), KEGG pathway classification (Kanehisa and Goto 2000; Kanehisa *et al.* 2016, 2017) and Saccharomyces Genome database (Cherry *et al.* 2012), we attributed each protein to a single functional category based on our expert knowledge (Table ST1). Considering the genes encoding the proteins, we also assigned to each protein a number of putative transcription factors (TFs). A total of 313 TFs with a consensus DNA-binding sequence were retrieved from the Yeastrack database (Teixeira *et al.* 2014; Abdulrehman *et al.* 2011; Monteiro *et al.* 2008; Teixeira *et al.* 2006).

### Statistical Methods

In order to estimate the genetic variance components for the different phenotypic traits, we adapted the model described in Lenarcic *et al.* (2012) to our particular half-diallel design that includes the diagonal with parental inbred strains from two species. Thus we included in our model intra- and inter-specific additive effects, inbreeding effects and intra- and inter-specific heterosis effects.

Formally, let *y_ijk_* be the observed phenotype for the cross between parents *i* and *j* in replica *k*. Our model reads:

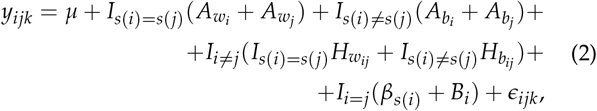

where:

- *μ* is the overall mean;
- *s*(*i*) associates to each parental strain *i* the specie it belongs to:

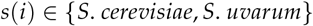
- 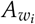 and 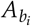 denote, respectively, the additive contributions of strain *i* in intra-specific (within species, *i*.*e*. *s*(*i*) = *s*(*j*)), and inter-specific (between species, *i*.*e*. *s*(*i*) ≠ *s*(*j*)) crosses;
- 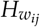 and 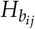 denote the interaction effect between parents (*i*, *j*) in intra-specific (within species) and inter-specific (between species) crosses, respectively. Due to our half-diallel design (no reciprocal crosses), they are assumed to be symmetric, *i*.*e*. 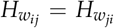 and 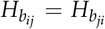. Hereafter we will refer to these effects as intra- and inter-specific heterosis effects, respectively;
- *β_s_*(*i*) and *B_i_* are, respectively, the deviation from the fixed overall effect for the species *s*(*i*) and the associated strain-specific contribution of strain *i* in the case of inbred lines. Hereafter we will refer to *Bi* as inbreeding effect;
- ϵ_*ijk*_ is the residual, the specific deviation of individual *ijk*;
- *I_condition_* is an indicator variable. Its value is equal to 1 if the condition is satisfied and 0 otherwise.

Therefore, for the parental lines we have:

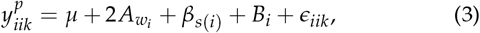

for the intra-specific hybrids:

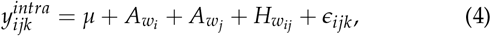

and for the inter-specific hybrids

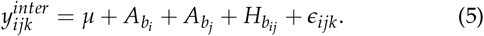

All genetic effects were considered as random variables drawn from a normal distribution. Formally, letting ***q*** ∈{***A_w_***, ***A_b_***, ***B***, ***H_w_***, ***H_b_}*** denote the genetic effect under consideration:

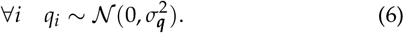

The full mixed-effect genetic model is thus defined by three fixed effects (the intercept *μ* and the inbreeding effects *β_Su_* and *β_Sc_*) and five genetic random effect variances 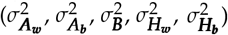

We did not declare mitochondrial effects because many genes encoding mitochondrial proteins are repressed under fermentation conditions, and because inter-specific hybrids harbor similar fermentation features for most fermentation kinetics and enological parameters whatever their mitochondrial genotype (Albertin *et al.* 2013a). In addition, we did not know the mitochondrial inheritance for most of the intra-specific crosses (table ST3).

### The fitting algorithm

Fixed effects, variance components of the genetic effects as well as their Best Linear Unbiased Predictors (*BLUPs*) were estimated using the *hglm* package in R (Ronnegard *et al.* 2010) that implements the estimation algorithm for hierarchical generalized linear models and allows fitting correlated random effects as well as random regression models by explicitly specifying the design matrices both for the fixed and random effects. The model, based on a maximum likelihood estimation, is deemed to produce unbiased statistics (Gumedze and Dunne 2011).

A separate analysis was conducted for each trait at each temperature, considering the vector of observations for the trait/temperature combination of interest, **y**, and re-writing model (eq. (2)) in matrix form:

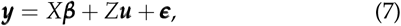

where X is the design matrix for the fixed effects, Z the design matrix for the random effect, ***β*=** (*μ,β_S.u._, β_S.c._*) and ***u*** = (***A_w_***, ***A_b_***, ***B***, ***H_w_***, ***H_b_***) are respectively the vectors of fixed effect parameters and random effect parameters, and ***ϵ*** is the vector of residual errors. With this notation, the construction of the model is straightforward from the data (for details see The fitting algorithm in Supplementary Materials).

Whenever the full model (eq. 2) failed to converge, we considered the subsequent model obtained by removing one effect at a time following the hierarchy imposed by the order of the fitting algorithm, *i.e.* first heterosis, second inbreeding effects and finally additive effects. The full model converged for all proteomic data. For the fermentation traits, the model did not converge for most of the *Ethyl* esters (*Ethyl-propanoate*, *Ethyl-butanoate*, *Ethyl-hexanoate*, *Ethyl-octanoate* and *Ethyl-decanoate*), as well as for *Acetate Ratio* and for *Acetic acid* that were removed from the analysis. For all other fermentation traits, the full model converged, except for *t.lag* at 18°, for which the additive model applied. For this trait, other genetic variance components were set to zero.

In order to test the robustness of the results, a bootstrap analysis was performed by sampling the 55 hybrids with replacement, conditionally to the 11 parental strains. Each bootstrap sample was submitted to the same analysis as described above. For each variance component, we checked that the estimations in the experimental sample were close to the median of the estimations in the bootstrap samples.

### Testing for the reliability of the model

Computer simulations were performed to test the statistical power of the *hglm* algorithm in predicting the values of the observables while producing unbiased estimations of the model parameters. We simulated a half-diallel between 11 strains, seven belonging to a species, four to the other. We computed the phenotypic values of each simulated cross by first drawing *μ*, *β_specie_*_1_, *β_specie_*_2_, 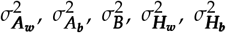 and 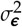 from a Gamma distribution fitted from the values estimated by the model on our dataset (see fig. SF1). Second, for each random effect ***q*** ∈{***A_w_***, ***A_b_***, ***B***, ***H_w_***, ***H_b_***, ϵ} we drew

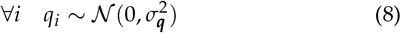

and computed the phenotypic values as in eq. 2, generating three replicas per cross.

We repeated the simulation 1000 times. We fitted the model and checked that the estimation of the random effects, the predicted phenotypic values as well as their variance components were close enough to the true values (fig. 1) and we noticed that inbreeding parameters were the most variable (fig. SF2 in *Supplementary figures*).

**Figure 1.**
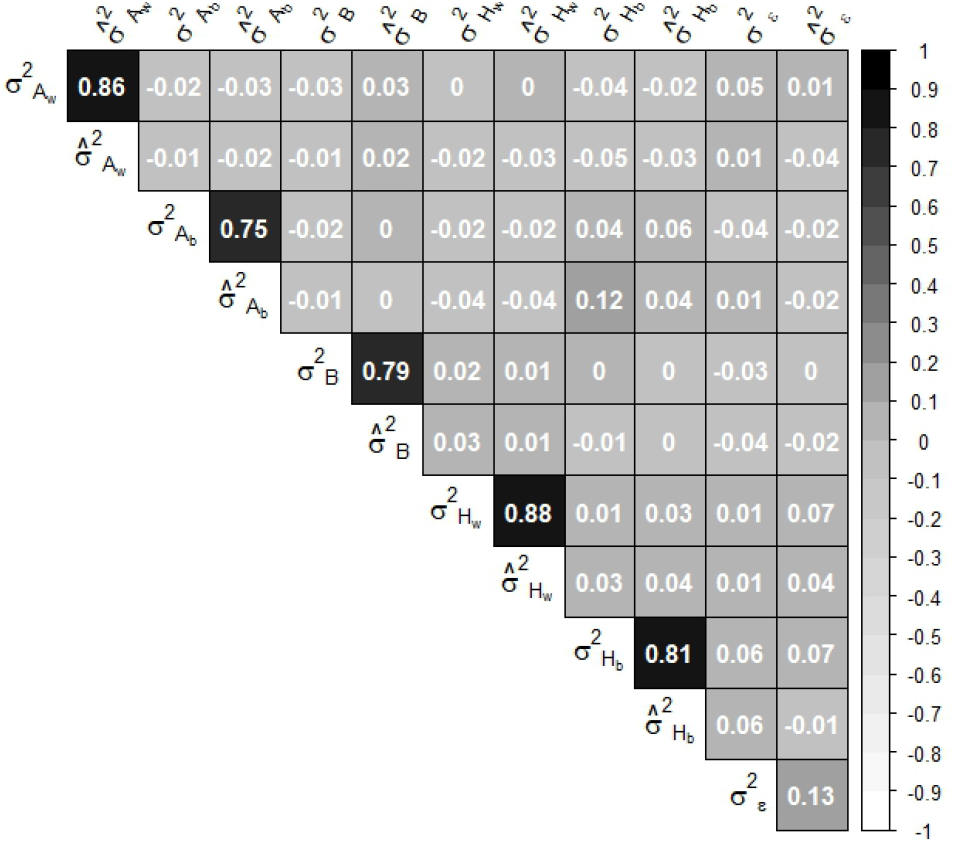
Correlation between estimated variance components and their true value. Variances have been estimated on a simulated half-diallel between 11 parental strain (seven belonging to a specie, four to the other). Phenotypic values have been computed as detailed in section Testing for the reliability of the model.

In addition, since we were interested in the correlation structure between the variance components of the genetic effects, we checked that possible correlations between random effects were not a statistical artifact of the model. Therefore, we simulated uncorrelated variances of random effects and we checked that no correlation structure was found between the estimated variance components, as can be seen in fig. 1. Simulations performed with different numbers of parental lines led to similar results (not shown).

### Fermentation traits

Before fitting our model, we updated eq. 2 in order to account for a block effect:

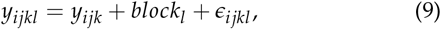

assuming that

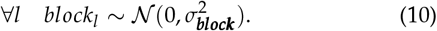

Many fermentation traits, mostly aromatic, were *log*-transformed in order to deal with the variable mean of the residuals. So as to handle the null values in the observations, we chose to consider the following transformation:

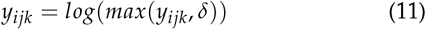

where *δ ~* 𝒰(0, *min*(***y***)). In this situation, as we introduced a random term in our analysis, which may skew parameter estimation, we decided to: (i) perform the *log*-transformation, (ii) compute the fitting algorithm, (iii) record the parameter’s estimation, then after having computed it a hundred times, (iv) consider the median of the estimators in order to achieve a more robust statistics.

### Protein abundances

For each cross, protein abundances have been quantified on average. Yet, to perform a diallel analysis at the proteomic level, replicas are critical for quantifying genetic variation. Therefore, we generated pseudo replicas using the residual variance estimated when quantifying protein abundances (Blein-Nicolas *et al.* 2013). Formally, let *y_ij_* be the average protein abundance of the cross between parents *i* and *j*. We generated three replicas as follows:

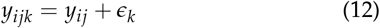

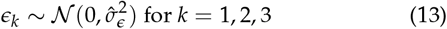

where 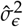 is the residual variance. Simulations of pseudo replicas and parameter estimations were performed 100 times. The final value of the parameters was the median of its estimation.

### Variance component analysis

For each trait, our mixed model generates a vector of variance components

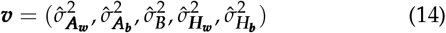

and the results were summarized in a matrix with rows being the different trait by temperature combinations, and columns the relative contribution of each component to the total genetic variance of the trait. We chose to perform unsupervised classification to compare the distributions of variance components between traits. Following the recommendations of Kurtz *et al.* (2015), percentages of variance components were transformed into real numbers using the following *clr*-transformation:

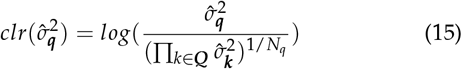

where *N_q_* is the total number of random effects and ***Q*** is the set of random variables fitted by the model. For fermentation traits, *N_q_* = 7 (accounting for block and residual variances, eq. 9), while *N_q_* = 6 for proteomic traits (eq. 2). We chose the *clr*-transformation because it satisfies *scale invariance*, *sub-compositional dominance* and *perturbation invariance* properties (Tsagris *et al.* 2011). Therefore the distance relationship between the original profiles is preserved by the selected sub-vectors thanks to the sub-compositional dominance property of the *clr*-transformation (see section Subcompositional dominance and distances in Supplementary Materials). The *clr*-transformation allowed us to test finite Gaussian mixture models using model-based clustering proposed in the *Mclust* package in R (Scrucca *et al.* 2016). Percentage of good assignments were computed by separating the data into training and validation sets.

This procedure was first applied separately for proteomic and fermentation traits (see Structuration of genetic variability at the fermentation trait level in Supplementary Materials). Protein groups were tested for enrichment in either Kegg pathways, transcription factors and heterotic proteins. Fermentation traits were tested for enrichment in the different trait categories (kinetic parameters, life-history, basic enological parameters, aromatic traits). For each cluster, Pearson’s chi-square test of enrichment was computed on protein functional category frequencies taking as prior probability the expected categorical frequency found in the MIPS database.

Further, fermentation traits were assigned to clusters identified on protein abundances profiles based on their membership probability computed through Gaussian finite mixture models.

### Data Availability

The data that support the findings of the current study are available from the corresponding author upon request. Supplementary materials contain:

- Demonstration of the relationship between the *subcompositional dominance* property and distances in the Euclidean space;
- Detailed description of the fitting algorithm;
- Description of the construction of the simulated values on a half-diallel design based on the genetic models supposed to explain heterosis and inbreeding;
- Demonstration of the equality between the variances of heterosis and inbreeding effects in three parents half-diallel designs with no maternal effects;
- Clustering analysis for the fermentation and life-history traits;
- Strains characterization based on the estimated BLUP of their genetic effects;
- Table ST1: Protein functional category classification (avail-able at figshare DOI:10.6084/m9.figshare.6683666);
- Table ST2: Raw values of genetic variances and broad sense heritability (BSH) estimated and analyzed in this study for protein abundances, and fermentation and life-history traits (available at figshare DOI:10.6084/m9.figshare.7128152);
- Table ST3: Mitochondrial inheritance of the phenotyped crosses of our study;
- Table ST4: Table of results from the Pearson’s chi-square test of cluster enrichment in proteins with a particular functional category;
- Figure SF1: Density distribution of the genetic variances estimated by the model;
- Figure SF2: Predicted BLUPs and phenotypic values versus their prior value used to compute the values of simulated diallels;
- Figure SF3: Clustering profiles of fermentation and life-history traits;
- Figure SF4: Global correlations of the genetic variance components for both protein abundances and the more integrated traits;
- Figure SF5: Representation of the standardized Pearson’s chi-square residuals of each cluster computed at 18° versus those at 26° estimated for the analysis of cluster enrichment in proteins with a particular functional category;
- Figure SF6: Correlation plot between genetic effects of fermentation and life-history trait profiles;
- Figure SF7: Intra-cluster correlations of variance components profiles for fermentation and life-history traits;
- Figure SF8: Variance components of fermentation and life-history traits at the two temperatures;
- Figure SF9: Summary example of the density distribution of a genetic variance estimation through bootstrap analysis;
- Figure SF10: Representation of the relationship between the variances of heterosis and inbreeding effects simulated through different genetic models;
- Figure SF11: For each trait and for each genetic effect are shown the strains with highest and lowest contribution at both temperatures;
- Figure SF12: For each trait are shown the estimated BLUPs of each genetic parameter.

## Results

In order to estimate genetic variance components from a diallel cross involving two yeast species, we proposed a decomposition of genetic effects based on the model of Lenarcic *et al.* (2012) that allowed to split the classical General (GCA) and Specific (SCA) Combining Abilities into intra- and inter-specific additive and heterosis effects, and to take into account inbreeding effects, defined as the difference between the inbred line value and the average value of all the crosses that have this inbred as parent.

Simulations showed that despite the small number of parents in the diallel, our model led to unbiased estimations of variance components, and that correlations between variance components did not arise from unidentifiability of some model’s parameter (fig. 1). Significance of variance components was assessed by bootstrap sampling. We found that whenever the fitting algorithm converged, variance component estimations were significant. For some traits and some variance components, the bootstrap distributions of the estimated variances were bimodal, suggesting a strong influence of a particular hybrid combination. However, the estimates were globally closed to the median of the bootstrap distribution (see example fig. SF9). Therefore, we are confident with our estimations, conditionally to the parents of the diallel.

Because temperature has a major effect on many traits and because, in previous work, numerous × strain temperature effects have been detected (da Silva *et al.* 2015; Blein-Nicolas *et al.* 2015), the model was applied to each trait separately at the two temperatures. We obtained estimations of fixed and random effect parameters, their corresponding variances, residuals and residual variances. For each trait, normality of residuals and homogeneity of variances was checked. Broad sense heritability was measured as the ratio of the sum of genetic variance components to the total phenotypic variance. It varied between 0.05 to 0.98 for protein abundances and between 0.04 to 0.95 for fermentation traits. Altogether, protein abundance measurements were highly repeatable (median heritability of 0.53), while fermentation traits were more variable. Median broad sense heritability was 0.77 for fermentation kinetic trait, 0.49 for life-history traits, 0.36 for basic enological products and 0.32 for aromatic traits. Whatever the amount of residual variance, all genetic variance components were significant for all traits, except for *t.lag* at 18°, for which only the variances of additive effects were significant. We found that variances associated to each genetic effect differ in a large extent between the two temperatures (shown for fermentation traits in fig. SF12).

Because of their potential interest for wine-making, BLUPs of fermentation traits are presented in section Strain characterization of Supplementary Materials. In the following, we focus on genetic variance components.

### Structuration of genetic variance components at the proteomic level

A Gaussian mixture model was used to classify the proteins according to their genetic variance components. The best model clearly identified nine clusters, each characterized by a particular profile of genetic variance components (fig. 2). Cluster 1 (88.4% of good assignments) consists of 11 proteins that have high variance of intra-specific heterosis effects and the smallest variance of inter-specific heterosis effects. Clusters 2, 4 and 9 have a very small variance of inbreeding effects. Clusters 2 and 4 differ from cluster 9 by their significant variance of inter-specific additive effects. 6.4% of proteins from cluster 2 (composed of 168 proteins with 93.2% of good assignments) can be attributed to cluster 4 and 10.4% of proteins from cluster 4 (65 proteins, 80.5% good assignments) to cluster 2. Proteins from clusters 3 (80.5% of good assignments) and 7 (93.3% of good assignments) have similar profiles.

**Figure 2.**
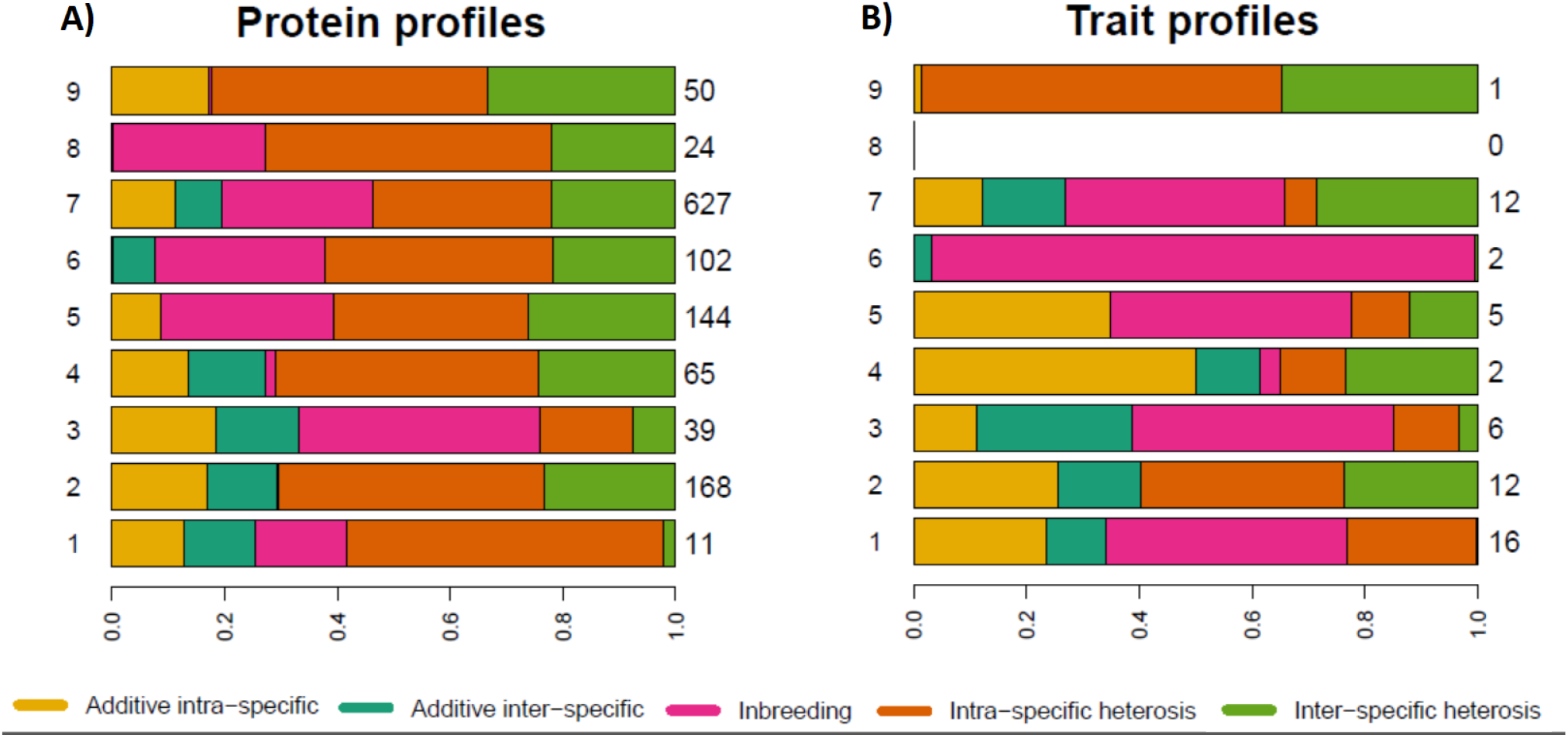
Clustering profiles of genetic variance components for protein abundances (A) against profiles of fermentation traits (B) predicted in each cluster. Cluster numbers are reported on the left, on the right the number of proteins or traits found in each cluster.

Indeed, 19.5% of the proteins from cluster 3 can be attributed to cluster 7 and 4% of the proteins from cluster 7 can be attributed to cluster 3. Cluster 3 consists in 39 proteins with relatively higher variance of additive and inbreeding effects. Cluster 7 has 627 proteins with higher variance of heterosis effects. Proteins from cluster 5 (144 proteins, 96% of good assignments) have significant variance of intra-specific additive effects but null variance of inter-specific additive effects and high heterosis and inbreeding effects variances. On the contrary, cluster 6 (102 proteins, 96.2% of good assignments) has null variance of intra-specific additive effects, small variance of additive inter-specific effects, and high variance of heterosis and inbreeding effects. Cluster 8 (96.9% of good assignments) consists of 24 proteins that have null variances of additive effects and high variances of heterosis and inbreeding effects. Finally, the 50 proteins in cluster 9 (95.4% of good assignments) are characterized by a null variance of additive inter-specific and inbreeding effects and high variance of intra-specific and inter-specific heterosis effects. Overall the same protein is generally found in two different clusters at the two temperatures (only 37% of proteins belong to the same cluster at the two temperatures).

The nine clusters were also clearly distinguishable from each other from their pattern of correlation between variance components (fig. 3). Globally, all variance components are negatively correlated, except for the variances of heterosis effects, 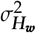 and 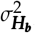 that are positively correlated (*r* = 0.47, fig. SF4).

**Figure 3.**
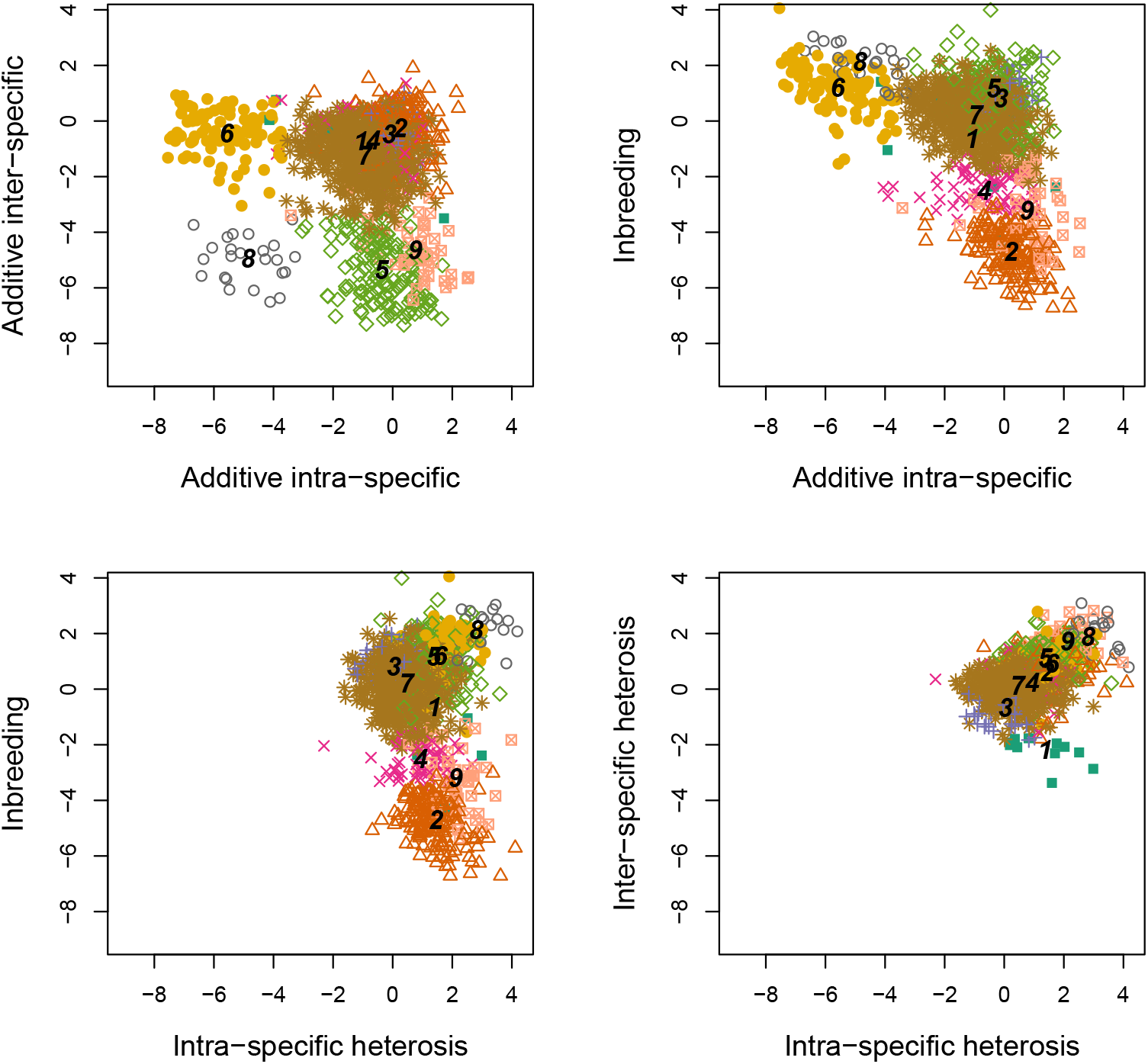
Patterns of correlations between genetic variance components of protein abundances. Points correspond to proteins, type and color combinations identify the clusters obtained by their classification based on a Gaussian Mixture model. Numbers from 1 to 9 identify class centers for each cluster.

Therefore, we can state that the 615 proteins at 18° and 26° form highly structured and well defined clusters according to their genetic variance component profiles.

### Proteins sharing a similar variance component structure share functional properties

In each protein cluster we tested for enrichment in functional categories at the two temperatures separately. Clusters were split into two groups of proteins, those measured at 18° and those measured at 26°, and the enrichment analysis was performed for each group. The statistical tests were significant for each cluster, except for cluster 1 at 18° and cluster 6 at 26° (tab. ST4). Even though one protein generally falls into two different clusters at two different temperatures, functional enrichments were globally the same at the two temperatures. Indeed, we found a high correlation between Pearson’s chi-squared residuals at both temperatures, except for clusters 3 and 9 (fig. SF5). Whenever a functional category was enriched/depleted at one temperature, it also tended to be enriched/depleted at the other temperature.

Cluster 1 is enriched with proteins quantified at 26° linked to response to stress, mating and transcription, and depleted with proteins related to cell fate and protein synthesis. Cluster 3 is enriched with proteins measured at 18° linked to amino-acid and nucleotide metabolism, and at 26° to cell fate and response to stress. Cluster 6 is enriched with proteins quantified at 18° linked to protein synthesis and nucleotide metabolism, and depleted in proteins linked to metabolism, other than amino acid, nucleotide and carbon metabolism. Cluster 9 is enriched in proteins linked to transcription at both temperatures, it is enriched in proteins measured at 18° linked to response to stress and mating, and depleted in proteins linked to protein synthesis and cell fate; at 26° it is enriched in proteins linked to nucleotide metabolism and transport. The other protein clusters have the same profile at both temperatures. Cluster 2 is enriched with proteins linked to amino-acids and carbon metabolism, cell fate and response to stress, and depleted in proteins linked to transport and mating. Cluster 4 is enriched in proteins linked to amino-acid metabolism, and to stress response at 26°. Cluster 5 is enriched in proteins linked to protein synthesis, amino-acid, nucleotide and other but not carbon metabolism, and depleted in proteins linked to transcription. Cluster 7 is enriched in proteins linked to amino-acids and carbon metabolism, and depleted in proteins linked to transcription, transport and signal. Cluster 8 is enriched in proteins linked to cell fate, stress response, nucleotide metabolism and mating, and depleted in proteins linked to other metabolisms, transport and protein synthesis. Hence, genetic variance components tend to cluster proteins having similar functions at both temperatures.

Concerning the number of transcription factors, we found no correlation between the number of transcription factors and the components of genetic variation of protein abundances.

Finally, Pearson’s chi-square test have been performed in order to investigate if there were differences between clusters regarding the proportion of heterotic proteins quantified in Blein-Nicolas *et al.* (2015). Results are shown in tab. 1: cluster 1, 2, 4 are enriched with heterotic proteins while in clusters 5, 7, 9 heterotic proteins are scarce (*χ*^2^ = 54.29, p-value<0.05). Hence, heterotic proteins are preferably found in clusters characterized by low variance of inbreeding effects and high variances of intra-specific and inter-specific heterosis effects.

**Table 1.**
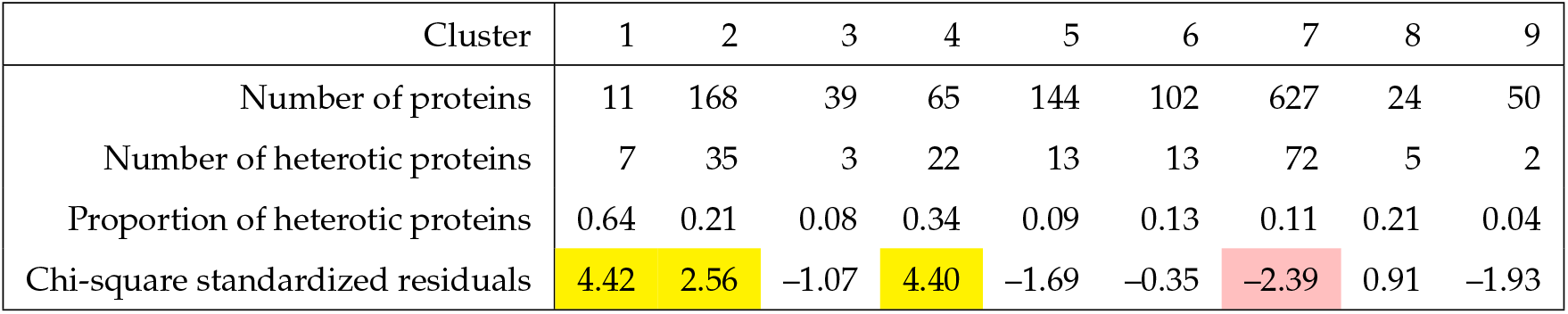
Pearson’s chi-square test for count data: comparison between the number of heterotic proteins in each cluster and group membership probability. The statistics clearly highlight clusters enriched of heterotic proteins (p-value<0.05). In yellow (respectively pink) are highlighted the clusters significantly enriched (resp. depleted) in heterotic proteins.

Briefly, despite poor correlations between variance components measured for the same protein at two temperatures, the nine clusters of proteins identified from the distribution of variance components group together proteins of similar function, based on their functional annotation. Heterotic proteins that show non-additive inheritance between parents and hybrids are mostly found in protein clusters with high variances of intra-specific and inter-specific heterosis effects and low variance of inbreeding effects.

### Variance components of fermentation traits fall into the proteomic landscape

Using for the fermentation/life history traits the same clustering approach as for the proteins, we clearly identified three profiles of genetic variance components (fig. SF3; see description in the section Structuration of genetic variability at the fermentation trait level of Supplementary Materials).

In order to compare the patterns of genetic variation of protein abundances and fermentation traits, we tried to assign fermentation traits to proteomic clusters based on the Gaussian Mixture model fitted on protein abundances profiles, as explained in section Variance component analysis of Materials and Methods. We chose for each fermentation trait the cluster of maximal membership probability. Most traits were assigned to a single protein cluster with a probability higher than 80%. The exceptions were *Sugar*/*EthanolYield* (26°), *X*4*MPP* (26°), *t*.75 (26°), *t*.*lag* (26°) and *t*.*lag* at both temperatures. Average variance components for each cluster are represented in fig. 2. Altogether, the 56 fermentation traits fall into eight proteomic clusters, most of them being assigned to clusters 1 (16 traits), 2 (12 traits), 7 (12 traits), 3 (6 traits), 5 (5 traits). Note that no trait was assigned to cluster 8, which corresponds to the cluster with the lowest variances of additive effects. Despite similarities with protein abundance traits, fermentation traits are characterized by higher variance of additive and inbreeding effects and globally higher contrasts in genetic variance components (fig. 4). Overall, 8 traits were attributed to the same cluster at the two temperatures: *J_max_*, *r*, *t-N_max_*, *Viability-t-75*, *X4MMP*, *Hexanoic acid*, *Hexanol*, *Ethanol*.

**Figure 4.**
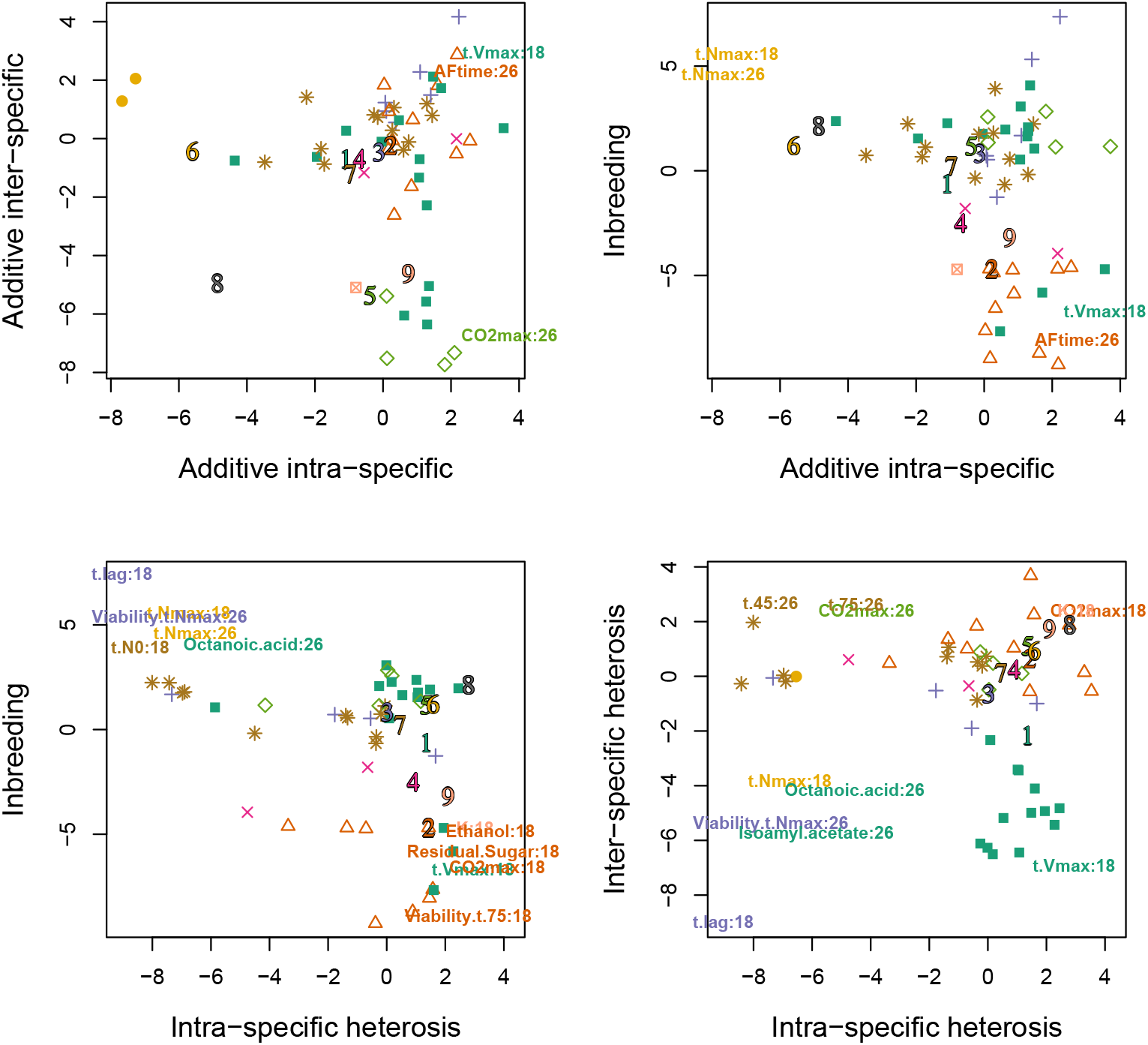
Variance components of fermentation traits. Fermentation traits are assigned to clusters identified at the proteomic level based on their membership probability computed through Gaussian finite mixture models. They are identified by the type and color combination of the cluster to which they are assigned. Numbers 1 to 9 identify class centers for each protein cluster. Labels are only given for outlier traits, *i.e.* those that do not belong to the 95% confidence interval of the genetic variance estimates of protein abundances on the plotted direction.

In addition, we investigated, for each temperature, the link between protein category in each cluster and type of fermentation trait. We see that at 18°, most Basic Enological Parameters (BEP) fall in cluster 2 where we found proteins involved in metabolism and stress response. Life History Traits fall in cluster 7 (amino-acid and carbon metabolism) and carrying capacity *K* falls in cluster 9 (cell growth) while *t-N_max_* is found in cluster 6 (nucleotide metabolism and protein synthesis). At 26°, most Aromatic Traits fall in cluster 1 (cell fate, stress response), most Fermentation Kinetics traits are found in cluster 7 (amino-acid and carbon metabolism), and BEP are in cluster 4 (stress response).

In conclusion, traits are generally attributed to different clusters at the two temperatures, based on the underlying components of genetic variation. Those clusters are characterized by the enrichment in proteins with a certain functional category, that may vary between temperatures. Interestingly, we found an association between traits linked to different metabolic processes and proteins involved in such processes just by taking into account their genetic variance decomposition.

### Intra-cluster correlations between variance components

Pearson’s correlation coefficients were computed for each pair of variance components within each cluster of proteins. Results clearly show different correlation structures between groups, particularly concerning correlation between the variances of heterosis and inbreeding effects (fig. 5). In cluster 1, variances of additive effects strongly and negatively correlate with each other. In cluster 3, there is a slightly negative correlation between 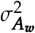 and the variances of heterosis effects, and there is a strong correlation between 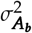 and variance of inbreeding effects. Cluster 4 is characterized by a weak negative correlation between 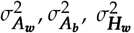 variances, and between 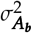, and the variances of heterosis and inbreeding effects. Clusters 5 and 7 preserve the global correlation structure. In cluster 2, the variances of intra-specific heterosis and inbreeding effects are negatively correlated, in cluster 6 the variances of heterosis and inbreeding effects are positively correlated, in cluster 8 the variances of inter-specific heterosis and inbreeding effects are positively correlated, and in cluster 9 the variances of heterosis and inbreeding effects are negatively correlated. Altogether, when a statistical significant correlation between the variances of additive, heterosis and inbreeding effects is found, it is negative.

**Figure 5.**
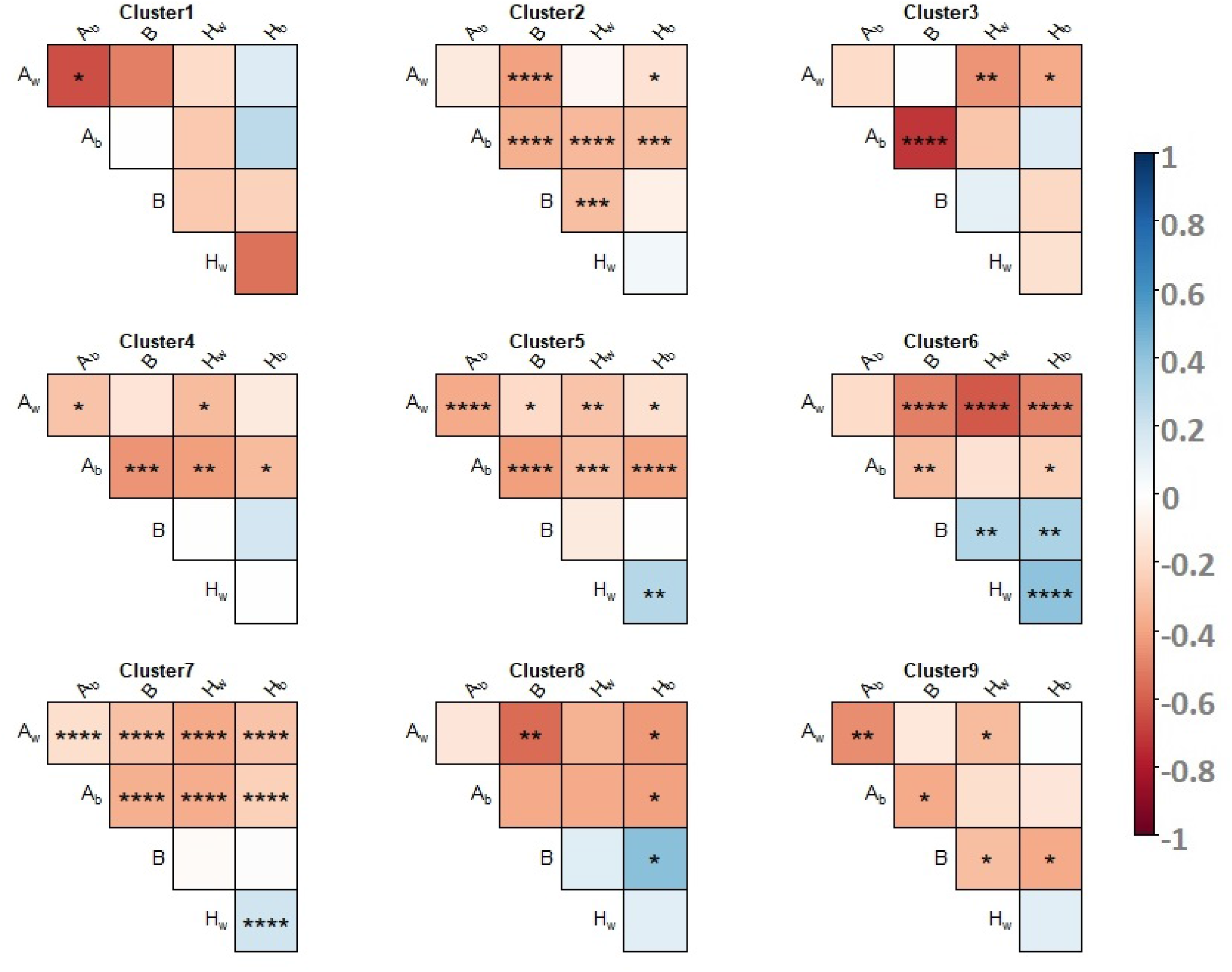
Pearson’s correlation test performed to investigate the intra-cluster correlations on proteomic data. For each cluster, correlation between variances of the genetic effects are indicated by a color-code. Warm colors stand for negative correlations and cold colors for positive correlations. * significant at *p* < 0.05; ** significant at *p* < 5 10^−3^; *** significant at *p* < 5 10^−4^; **** significant at *p* < 5 10^−5^. No symbol: not significant.

Variances of additive effects tend to be negatively correlated to variances of heterosis and inbreeding effects, and there is no straightforward relationship between the variances of heterosis and inbreeding effects: 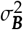 can be either negatively (cluster 9) or positively (cluster 6) correlated to both 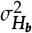 and 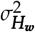, negatively correlated to 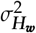 (cluster 2), positively correlated to 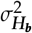 (cluster 8). However, 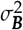 can also be independent from either 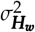 or 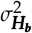 (cluster 1,2,3,4,5,7,8).

## Discussion

In this paper, we focused on the comparative analysis of genetic variance components estimated through the decomposition of traits value quantified in a half-diallel cross during or at the end of alcoholic fermentation. The cross design involved 11 yeast strains from two related species naturally associated with wine fermentations, *S. cerevisiae* and *S. uvarum*, and the set of traits quantified spanned from protein abundances to fermentation and life-history.

Genetic variances have been estimated through a comprehensive genetic model that allowed us to decompose the phenotypic value of a cross, including the parental inbred strains, in terms of additive and interaction effects. This decomposition can be described in the following way. The parental inbred lines have two identical haploid genomes, while the hybrids have two different haploid genomes, each inherited by one parent. Additive effects refer to the average value conferred by a single haploid genome with respect to any other haploid genome, and interaction effects refer to the non-additive effect of a particular genotype computed as the difference between the particular diploid value and the average additive effect of its haploid genomes. The presence of the parental inbreds in the experimental design permits a decomposition of those effects into heterosis and inbreeding effects. Inbreeding effect is defined as the difference between the value of the inbred strain (with the same haploid genome twice) and the average of all the crosses having at least one copy of the haploid parental genome. Heterosis effect is defined as the difference between a single pairwise genome combination and the average value of hybrids having one or the other haploid genome. Thanks to the presence of two different yeast species in our experimental design, we could distinguish intra-specific and inter-specific genetic effects. Indeed, the additive effect of a strain and the heterosis effect of a hybrid between two strains may differ depending on whether the strains belong to the same species or not. Therefore, intra-specific (respectively inter-specific) additive effect refers to the average value conferred by a single haploid genome with respect to any other haploid genome from the same specie (respectively from another species), and intra-specific (respectively inter-specific) heterosis effect refers to the difference between a single pairwise genome combination from the same specie (respectively from the two species) and the average value of the intra-specific (respectively inter-specific) hybrids having one or the other haploid genome.

This general model could be adapted to consider mitochondrial effects, which we did not declare for biological and technical reasons given in Materials and Methods. If such effects do exist in our genetic material they are expected to be weak and confounded with other effects.

The variance components of the genetic effects defined above have been estimated using the linear mixed model (*LMM*) described in eq. 2. Whenever a variance component was significant, it meant that genetic differences were found between strains. We checked the ability of the *LMM* to estimate genetic parameters by means of computer simulations and the robustness of the estimations through bootstrap analysis. In the simulations, despite residual variances that were not well correlated to their true value, estimated genetic variances were found to highly correlate with their true value (fig. 1). However, residuals quantified on the proteomic data highly correlate with their true value (see section Protein abundances). Bootstrap analysis, performed by sampling the 55 hybrids with replacement, conditionally to the 11 parental strains, revealed that for each variance component the estimations in the experimental sample were close to the median of the estimations in the bootstrap samples. For some traits and some variance components, the distribution of the bootstrap estimated variances were bimodal, suggesting a strong influence from a particular hybrid combination. However, it was never flat or smooth, in agreement with the non arbitrary choice of the parameters. Therefore, we are confident about the estimations of the genetic variances, conditionally to the parents of the diallel.

We were able to characterize the 615 proteins and the 28 fermentation and life-history traits quantified at 18° and 26° by a particular profile of genetic variance components despite the small number of parental inbred strains from which the half-diallel was built. We found that variances of intra- and inter-specific effects differed in a large extent, pointing out that the genetic effects are highly influenced by crossing strains from the same species or not. The degree of intra- and inter-specific genetic variation captures the evolutionary history the two species have undergone for the different traits. For instance, traits with a low variance of intra-specific additive effects but high variance of inter-specific additive effects have a high potential to evolve in inter- but not intra-specific crosses.

Each trait has been treated at each temperature separately, considering trait × temperature as independent characters. Indeed, genotype-by-environment interactions affect very commonly phenotypic variation. In particular, it is well documented that the genetic architecture of a trait is not stable under varying environments, highlighting the fact that evolutionary processes may depend largely upon ecological conditions (Falconer 1960; Lynch and Walsh 1998; Hermisson and Wagner 2004; Robinson *et al.* 2009; Malosetti *et al.* 2013). Accordingly we found a weak correlation between genetic variances at the two temperatures.

The molecular phenotypes (protein abundances) reflect the underlying genetic factors involved in the cellular processes regulating the most integrated traits. So we investigated the distribution of the components of genetic variation of protein abundances in relation to fermentation and life-history trait variance components. We found nine clear-cut clusters of protein variance components, and we were able to assign traits to these clusters based on their genetic variance components. Overall, the profiles of the fermentation and life-history traits associated to each cluster were close to that of the proteomic level, but they were characterized by higher variance of additive effects; further, we could not assign any trait to cluster 8, which has null variance of additive effects, *i.e.* which is the group with the less heritable proteins. Altogether these results reveal that the most integrated traits have a higher evolutionary potential compared to protein abundances.

We tested for cluster enrichment in protein functions, based on the functional annotation of the proteins. Clusters were found to group together proteins of similar functions. Despite the fact that 63% of the proteins were found in different clusters at the two temperatures, the metabolic functions were preserved. This suggests temperature-specific regulatory changes that achieve the maintenance of cell functions. At the trait level, 16 over 28 fermentation/life-history traits (57%) fell into the same cluster at the two temperatures (fig. SF8). For the 12 remaining traits, changes in the distribution of variance components between the two temperatures can be explained by *G × E* interactions.

Beside, we have shown that the clusters were characterized by a particular profile of genetic variance components, which suggests that traits that group together share a similar evolutionary history. If all traits were neutral, they would have shown the same equilibrium level of total genetic variance of approximately 2*NV_m_* (*N* the effective population size and *V_m_* the mutational variance (Lynch and Hill 1986)) with a similar partition of genetic variance components. The existence of different profiles of variance components probably reflects that the different types of traits have been subject to particular selective pressures.

Beyond, the nine clusters were clearly distinguishable from each other from their pattern of correlation between variance components. Overall, the variances of intra- and inter-specific additive effects were negatively correlated to the variances of heterosis and inbreeding effects. This may reveal differences in the patterns of allele frequencies at the underlying loci. In a biallelic case, additive genetic variance is always maximum for intermediate allele frequencies, while dominance and epistatic variances (which are components of the variances of heterosis and inbreeding effects) are maximum for more extreme allele frequencies (Hill *et al.* (2008)). A trait with a high variance of additive effects is therefore expected to have lower dominance or epistatic variances. Conversely, a trait with low variance of additive effects may exhibit high dominance and epistatic variances.

In the common view, heterosis and inbreeding are corollary effects. However, we have shown that the variances of heterosis and inbreeding effects could be negatively, positively or not correlated to each other. For a better understanding of such a decoupling, we simulated a half-diallel design between *N* parental strains (for details see section Half-diallel simulation construction in Supplementary Materials). We computed the phenotypic values of the parental lines and hybrids starting with a simple additive model (neither dominance at any locus nor epistasis), then we added dominance and/or epistasis effects. We considered different degrees of dominance for each couple of alleles (including dominance of the strongest allele, h=0) and *additive × additive* and *dominance × dominance* epistasis, and we let the number of alleles per locus to vary. We considered all possible combinations of these effects. Finally we decomposed the values of the simulated traits into additive, heterosis and inbreeding effects.

Not surprisingly, the variances of heterosis and inbreeding effects are both null when there is neither dominance nor epistasis. If there is *additive × additive* epistasis with no dominance, the variances of heterosis and inbreeding effects are strictly correlated, with very low variance of heterosis effects. In the other conditions, the results depend on the number of parental lines. With three parents, the variance of heterosis and inbreeding effects are strictly equal, as it can be shown analytically (see section Inbreeding depression and heterosis variances are equal in three-parent diallel in Supplementary Materials). Otherwise the correlation between the variances of heterosis and inbreeding effects varies in function of the number of loci affecting the trait of interest, on the frequency of alleles in the population and on the presence of dominance and epistatic effects. In general, the correlation between the variances of heterosis and inbreeding effects tends to become null when the number of parental lines, the number of alleles per locus and the number of loci increase. Given these parameters, whether there is dominance or not, and whatever the type of dominance, the lowest correlations between the variances of heterosis and inbreeding effects are observed when there are both types of epistasis together (fig. 6 and fig. SF10). However in no case we get negative correlations between the two variances. Further, we decided to consider the data obtained on all the different cases together and we run as previously a Gaussian Mixture Model to cluster genetic variances components. We computed intra-cluster correlations varying the number of alleles per locus, the number of loci and the distribution in which we drew allele values. Those correlations did not show profiles similar to those obtained with real data (correlations between genetic effects are commonly positive or null).

**Figure 6.**
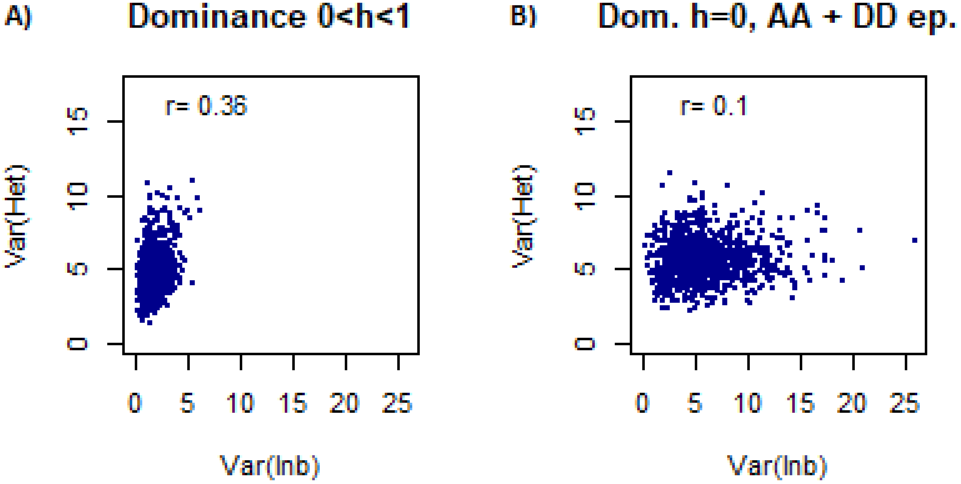
Correlation between the variances of heterosis and inbreeding effects for: A) Additive model with symmetrical dominance (no epistasis), B) Additive model with dominance of the strongest allele, additive ×additive and dominance ×dominance epistasis. The simulated half-diallel consisted of 11 parental lines. Phenotypic values were supposed to depend on 10 loci, and the number of alleles per loci was imposed to 11. Allele values were drawn from a gamma distribution (*k*=10, *θ*=20) and epistatic effects from a normal distribution (*𝒩* (0, 3)).

Classical genetic studies and modern molecular evolutionary approaches now suggest that inbreeding effects and heterosis are predominantly caused by the presence of recessive deleterious mutations in the population (Charlesworth and Charlesworth 1999; Charlesworth and Willis 2009). Therefore understanding the effects of selection against deleterious alleles is crucial. Population structure also plays a key role in this framework. Indeed, population subdivision increases homozygosity through inbreeding, an effective process for purging deleterious alleles, but it also decreases selection efficiency by decreasing the genetic diversity. Allele frequency changes also modify the genetic variance components (Hill *et al.* 2008; Barton 2017). A more complex model, which takes into account selection, allele frequency, population structure and the presence of deleterious mutations is thus needed to explain our observations. Glémin *et al.* (2003) have discussed about the patterns of correlation between inbreeding effects and heterosis in a structured population assuming low frequencies of deleterious mutations, only present in the heterozygous state. They defined within- and between-demes inbreeding depression as the decline in mean fitness of selfed individuals relative to out-crossed individuals within the demes and as the decline in mean fitness of selfed individuals relative to out-crossed individuals between demes, respectively; and heterosis as the excess in mean fitness of individuals produced by out-crosses between demes relative to mean fitness of individuals produced by out-crosses within the demes. They stated that population structure decreases within-demes inbreeding depression while it increases between-deme inbreeding depression, and that increasing the inbreeding coefficient reduces within- and between-deme inbreeding depression and heterosis. A similar result was obtained by Roze and Rous-set (2004) who considered a diffusion model in a population of partially selfing individuals subdivided according to an island model, with a large but finite number of demes. They found that generally within-deme inbreeding depression and heterosis are positively correlated upon selfing and, when the degree of population subdivision is high, inbreeding depression and heterosis are negatively correlated. To our knowledge, the present study reports the first experimental example of such a decoupling.

In conclusion, our findings have special relevance in three main directions: *(i) Detection of Quantitative Trait Loci (QTL).* Variances of additive effects are crucial for the detection of genes with significant quantitative effect, and variances of heterosis/inbreeding effects for the detection of gene-gene interactions when the part of genetic variance they explain is large; *(ii) Integration of proteomic data into Genome Scale Metabolic (GSM) model:* we assigned fermentation traits to clusters obtained on the components of genetic variation of protein abundances. Traits associated to a metabolic process were linked to proteins involved to such process, therefore we are confident that integrating proteins related to the most integrated traits into a GSM could improve their prediction, with particular attention to the prediction of heterosis; *(iii) Model heterosis and inbreeding variation:* we have highlighted various patterns of variation between the variances of heterosis and inbreeding effects that cannot be explained with simple quantitative genetics models. It would be interesting to construct *in silico* experiments to search for the key parameters that drive these patterns.

## Acknowledgments

We thank very much Dr. Arnaud Le Rouzic for exciting discussions and its material to purse the preliminary analysis of the diallel design. We thank very much Dr. Monique Bolotin for her help in the functional annotation of the proteins, and Dr. Warren Albertin and Dr. Philippe Marullo for their advice regarding yeast genetic material. This work was supported by a public PhD grant of the French National research Agency (ANR) as part of the "Investissement d’Avenir" program, through the “IDI 2015” project funded by the IDEX Paris-Saclay, ANR-11-IDEX-0003-02.

## Literature Cited

Abdulrehman, D., P. T. Monteiro, M. C. Teixeira, N. P. Mira, A. B. Lourenço, et al., 2011 Yeastract: providing a programmatic access to curated transcriptional regulatory associations in saccharomyces cerevisiae through a web services interface. Nucleic Acids Research 39: D136–D140.

Albertin, W., T. da Silva, M. Rigoulet, B. Salin, I. Masneuf-Pomarede, et al., 2013a The mitochondrial genome impacts respiration but not fermentation in interspecific saccharomyces hybrids. PLOS ONE 8: 1–14.

Albertin, W., P. Marullo, M. Aigle, A. Bourgais, M. Bely, et al., 2009 Evidence for autotetraploidy associated with reproductive isolation in saccharomyces cerevisiae: towards a new domesticated species. Journal of Evolutionary Biology 22: 2157–2170.

Albertin, W., P. Marullo, M. Bely, M. Aigle, A. Bourgais, et al., 2013b Linking Post-Translational Modifications and Variation of Phenotypic Traits. Molecular & Cellular Proteomics 12: 720–735.

Barton, N. H., 2017 How does epistasis influence the response to selection? Heredity (Edinb) 118.

Blein-Nicolas, M., W. Albertin, T., da Silva, B. Valot, T. Balliau, et al., 2015 A systems approach to elucidate heterosis of protein abundances in yeast. Mol Cell Proteomics 14: 2056–71.

Blein-Nicolas, M., W. Albertin, B. Valot, P. Marullo, D. Sicard, et al., 2013 Yeast proteome variations reveal different adaptive responses to grape must fermentation. Molecular Biology and Evolution 30: 1368.

Blein-Nicolas, M., H. Xu, D., de Vienne, C. Giraud, S. Huet, et al., 2012 Including shared peptides for estimating protein abundances: A significant improvement for quantitative proteomics. PROTEOMICS 12: 2797–2801.

Bulmer, M. G., 1980 The mathematical theory of quantitative genetics/ M.G. Bulmer. Clarendon Press; New York: Oxford University Press Oxford.

Charlesworth, B. and D. Charlesworth, 1999 The genetic basis of inbreeding depression. Genetical Research 74: 329–340.

Charlesworth, B., M. T. Morgan, and D. Charlesworth, 1991 Multilocus models of inbreeding depression with synergistic selection and partial self-fertilization. Genetics Research 57: 177–194.

Charlesworth, D. and J. H. Willis, 2009 The genetics of inbreeding depression. Nature Reviews Genetics 10: 783–796.

Cherry, J. M., E. L. Hong, C. Amundsen, R. Balakrishnan, G. Binkley, et al., 2012 Saccharomyces Genome Database: the genomics resource of budding yeast. Nucleic Acids Research 40: D700–705.

Cockerham, C. C. and B. S. Weir, 1977 Quadratic analyses of reciprocal crosses. Biometrics 33: 187–203.

da Silva, T., W. Albertin, C. Dillmann, M. Bely, S. la Guerche, et al., 2015 Hybridization within saccharomyces genus results in homoeostasis and phenotypic novelty in winemaking conditions. PLOS ONE 10: 1–24.

Davenport, C. B., 1908 Degeneration, albinism and inbreeding. Science 28: 454–455, WOS:000201859500057.

Eberhart, S. A. and C. O. Gardner, 1966 A General Model for Genetic Effects. Biometrics 22: 864–881.

Falconer, D. S., 1960 Introduction to quantitative genetics. New York,: Ronald Press Co.

Fievet, J. B., C. Dillmann, and D., de Vienne, 2010 Systemic properties of metabolic networks lead to an epistasis-based model for heterosis. Theoretical and Applied Genetics 120: 463–473, WOS:000272803700025.

Fiévet, J. B., T. Nidelet, C. Dillmann, and D. de Vienne, 2018 Heterosis is a systemic property emerging from nonlinear genotype-phenotype relationships: evidence from in vitro genetics and computer simulations. Frontiers in Genetics 9.

Glémin, S., J. Ronfort, and T. Bataillon, 2003 Patterns of inbreeding depression and architecture of the load in subdivided populations. Genetics 165: 2193–2212.

Gowen, J. W., 1952 Heterosis. Iowa state pr edition.

Graham, G. I., D. W. Wolff, and C. W. Stuber, 1997 Characterization of a yield quantitative trait locus on chromosome five of maize by fine mapping. Crop Sci. 37: 1601–1610, WOS:A1997XZ35500033.

Greenberg, A. J., S. R. Hackett, L. G. Harshman, and A. G. Clark, 2010 A hierarchical bayesian model for a novel sparse partial diallel crossing design. Genetics.

Griffing, B., 1956 Concept of general and specific combining ability in relation to diallel crossing systems. Australian Journal of Biological Sciences 9: 463–493.

Gumedze, F. and T. Dunne, 2011 Parameter estimation and inference in the linear mixed model. Linear Algebra and its Applications 435: 1920–1944.

Hallauer, A., W. Russell, and K. Lamkey, 1988 Corn breeding: 463–564. Sprague, GF and. IW Dudley: Corn and corn improvement. Agron. Monogr. (third edition). ASA, CSSA and SSSA Madison, WI.

Hallauer, A. R. and J. B. M. Filho, 1988 Quantitative Genetics in Maize Breeding. Iowa State University Press.

Hermisson, J. and G. P. Wagner, 2004 The Population Genetic Theory of Hidden Variation and Genetic Robustness. Genetics 168: 2271–2284.

Hill, W. G., M. E. Goddard, and P. M. Visscher, 2008 Data and theory point to mainly additive genetic variance for complex traits. PLOS Genetics 4: 1–10.

Huang, X., S. Yang, J. Gong, Q. Zhao, Q. Feng, et al., 2016 Genomic architecture of heterosis for yield traits in rice. Nature 537: 629–633.

Hull, F., 1946 Overdominance and Corn Breeding Where Hybrid Seed Is Not Feasible. Journal of the American Society of Agronomy 38: 1100–1103, WOS:A1946UC58500007.

Kanehisa, M., M. Furumichi, M. Tanabe, Y. Sato, and K. Morishima, 2017 KEGG: new perspectives on genomes, pathways, diseases and drugs. Nucleic Acids Research 45: D353–D361.

Kanehisa, M. and S. Goto, 2000 KEGG: kyoto encyclopedia of genes and genomes. Nucleic Acids Research 28: 27–30.

Kanehisa, M., Y. Sato, M. Kawashima, M. Furumichi, and M. Tanabe, 2016 KEGG as a reference resource for gene and protein annotation. Nucleic Acids Research 44: D457–462.

Kurtz, Z. D., C. L. Müller, E. R. Miraldi, D. R. Littman, M. J. Blaser, et al., 2015 Sparse and Compositionally Robust Inference of Microbial Ecological Networks. PLOS Computational Biology 11: e1004226.

Lande, R. and D. W. Schemske, 1985 The evolution of self-fertilization and inbreeding depression in plants. i. genetic models. Evolution 39: 24–40.

Lariepe, A., B. Mangin, S. Jasson, V. Combes, F. Dumas, et al., 2012 The Genetic Basis of Heterosis: Multiparental Quantitative Trait Loci Mapping Reveals Contrasted Levels of Apparent Overdominance Among Traits of Agronomical Interest in Maize (Zea mays L.). Genetics 190: 795–U835, WOS:000300621200037.

Lenarcic, A. B., K. L. Svenson, G. A. Churchill, and W. Valdar, 2012 A general bayesian approach to analyzing diallel crosses of inbred strains. Genetics 190: 413–435.

Lynch, M. and W. G. Hill, 1986 Phenotypic Evolution by neutral mutation. Evolution; International Journal of Organic Evolution 40: 915–935.

Lynch, M. and B. Walsh, 1998 Genetics and Analysis of Quantitative Traits. Sinauer, Google-Books-ID: UhCCQgAACAAJ.

Malosetti, M., J.-M. Ribaut, and F. A. van Eeuwijk, 2013 The statistical analysis of multi-environment data: modeling genotype-by-environment interaction and its genetic basis. Frontiers in Physiology 4.

Martì-Raga, M., E. Peltier, A. Mas, G. Beltran, and P. Marullo, 2017 Genetic Causes of Phenotypic Adaptation to the Second Fermentation of Sparkling Wines in Saccharomyces cerevisiae. G3: Genes, Genomes, Genetics 7: 399–412.

Melchinger, A. E. and R. K. Gumber, 1998 Overview of Heterosis and Heterotic Groups in Agronomic Crops. In Concepts and Breeding of Heterosis in Crop Plants, edited by K. R. Larnkey and J. E. Staub, pp. 29–44, Crop Science Society of America, USA.

Monteiro, P. T., N. D. Mendes, M. C. Teixeira, S. d’Orey, S. Tenreiro, et al., 2008 Yeastract-discoverer: new tools to improve the analysis of transcriptional regulatory associations in saccharomyces cerevisiae. Nucleic Acids Research 36: D132–D136.

Omholt, S. W., E. Plahte, L. Øyehaug, and K. Xiang, 2000 Gene Regulatory Networks Generating the Phenomena of Additivity, Dominance and Epistasis. Genetics 155: 969–980.

Powers, L., 1944 An expansion of Jones’s theory for the explanation of heterosis. The American Naturalist 78: 275–280.

Ramya, A. R., L. Ahamed M, C. T. Satyavathi, A. Rathore, P. Katiyar, et al., 2018 Towards Defining Heterotic Gene Pools in Pearl Millet [Pennisetum glaucum (L.) R. Br.]. Frontiers in Plant Science 8.

Redden, R., 1991 The effect of epistasis on chromosome mapping of quantitative characters in wheat. I. Time to spike emergence. Australian Journal of Agricultural Research 42: 1.

Robinson, M. R., A. J. Wilson, J. G. Pilkington, T. H. Clutton-Brock, J. M. Pemberton, et al., 2009 The Impact of Environmental Heterogeneity on Genetic Architecture in a Wild Population of Soay Sheep. Genetics 181: 1639–1648.

Ronnegard, L., X. Shen, and M. Alam, 2010 hglm: A package for fitting hierarchical generalized linear models. The R Journal 2: 20–28.

Roze, D. and F. Rousset, 2004 Joint effects of self-fertilization and population structure on mutation load, inbreeding depression and heterosis. Genetics 167: 1001–1015.

Ruepp, A., A. Zollner, D. Maier, K. Albermann, J. Hani, et al., 2004 The FunCat, a functional annotation scheme for systematic classification of proteins from whole genomes. Nucleic Acids Research 32: 5539–5545.

Schnable, P. S. and N. M. Springer, 2013 Progress toward understanding heterosis in crop plants. Annu Rev Plant Biol 64: 71–88.

Scrucca, L., M. Fop, T. B. Murphy, and A. E. Raftery, 2016 mclust 5: clustering, classification and density estimation using Gaussian finite mixture models. The R Journal 8: 205–233.

Seymour, D. K., E. Chae, D. G. Grimm, C. Martín Pizarro, A. Habring-Müller, et al., 2016 Genetic architecture of non-additive inheritance in Arabidopsis thaliana hybrids. Proc. Natl. Acad. Sci. U.S.A. 113: E7317–E7326.

Shao, H., L. C. Burrage, D. S. Sinasac, A. E. Hill, S. R. Ernest, et al., 2008 Genetic architecture of complex traits: Large phenotypic effects and pervasive epistasis. Proceedings of the National Academy of Sciences 105: 19910–19914.

Shull, G. H., 1908 The Composition of a Field of Maize. Journal of Heredity os-4: 296–301.

Sprague, G. F. and E. L. Tatum, 1942 General vs. specific combining ability in single crosses of corn. PROTEOMICS 34: 923–932.

Teixeira, M. C., P. Monteiro, P. Jain, S. Tenreiro, A. R. Fernandes, et al., 2006 The yeastract database: a tool for the analysis of transcription regulatory associations in saccharomyces cerevisiae. Nucleic Acids Research 34: D446–D451.

Teixeira, M. C., P. T. Monteiro, J. F. Guerreiro, J. P. Gonçalves, N. P. Mira, et al., 2014 The yeastract database: an upgraded information system for the analysis of gene and genomic transcription regulation in saccharomyces cerevisiae. Nucleic Acids Research 42: D161–D166.

Tsagris, M. T., S. Preston, and A. T. A. Wood, 2011 A data-based power transformation for compositional data. ArXiv e-prints.

Wright, S., 1934 Physiological and evolutionary theories of dominance. American Naturalist 68: 24–53, WOS:000200907000002.

Xiao, J., J. Li, L. Yuan, and S. Tanksley, 1995 Dominance Is the Major Genetic-Basis of Heterosis in Rice as Revealed by Qtl Analysis Using Molecular Markers. Genetics 140: 745–754, WOS:A1995RA36600028.

Zhu, J. and B. S. Weir, 1996 Mixed model approaches for diallel analysis based on a bio-model. Genetical Research 68: 233–240.

